# Assessment of environmental variables for species distribution modelling: insights from the mosaic distribution of red- and yellow-bellied toads

**DOI:** 10.1101/2025.04.24.650429

**Authors:** Jan W. Arntzen, Krisztián Harmos, Judit Vörös

**Affiliations:** Institute of Biology, Leiden University, Leiden, The Netherlands; Naturalis Biodiversity Center, Leiden, The Netherlands; Bükk National Park Directorate, 3304 Eger, Sánc utca 6, Hungary; HUN-REN Balaton Limnological Research Institute, 8237 Tihany, Klebersberg Kuno u. 3, Hungary

**Keywords:** *Bombina variegata*, Enclaves, Parameter selection, Parapatry, Transferability, Two-species distribution modelling

## Abstract

Species distribution modelling can possibly be improved through the preferential use of explanatory variables that reflect the natural history features of the species being modelled. Red- and yellow-bellied toads (genus *Bombina*) engage in an intricate mosaic distribution across Europe. Analysing new atlas data on these species’ mutual distribution in Hungary with principal coordinate analysis we identified their differential ecological preferences as forested, hilly and mountainous for *B. variegata* and open lowland for *B. bombina*. These locally operating parameters we consider to be good proxies for the essential species difference which resides in breeding in ephemeral puddles (*B. variegata*) versus larger permanent ponds (*B. bombina*). With two-species distribution modelling *–* in which the presence of one species is contrasted with the presence of the counterpart species *–* we obtained excellent model fit (AUC) for climate and elevation / land cover datasets alike (AUC=0.98 versus 0.95). For both models fit values dropped upon transference to surrounding countries, yet the latter model kept significantly higher predictive power (AUC=0.91) than the climate model (AUC=0.79). Swapping elevation for ‘hilliness’ as suggested in the literature had a significant negative effect on model performance. We conclude that an informed parameter selection enhances model transferability, therewith improving our understanding of species-habitat associations.

**Graphical abstract:** 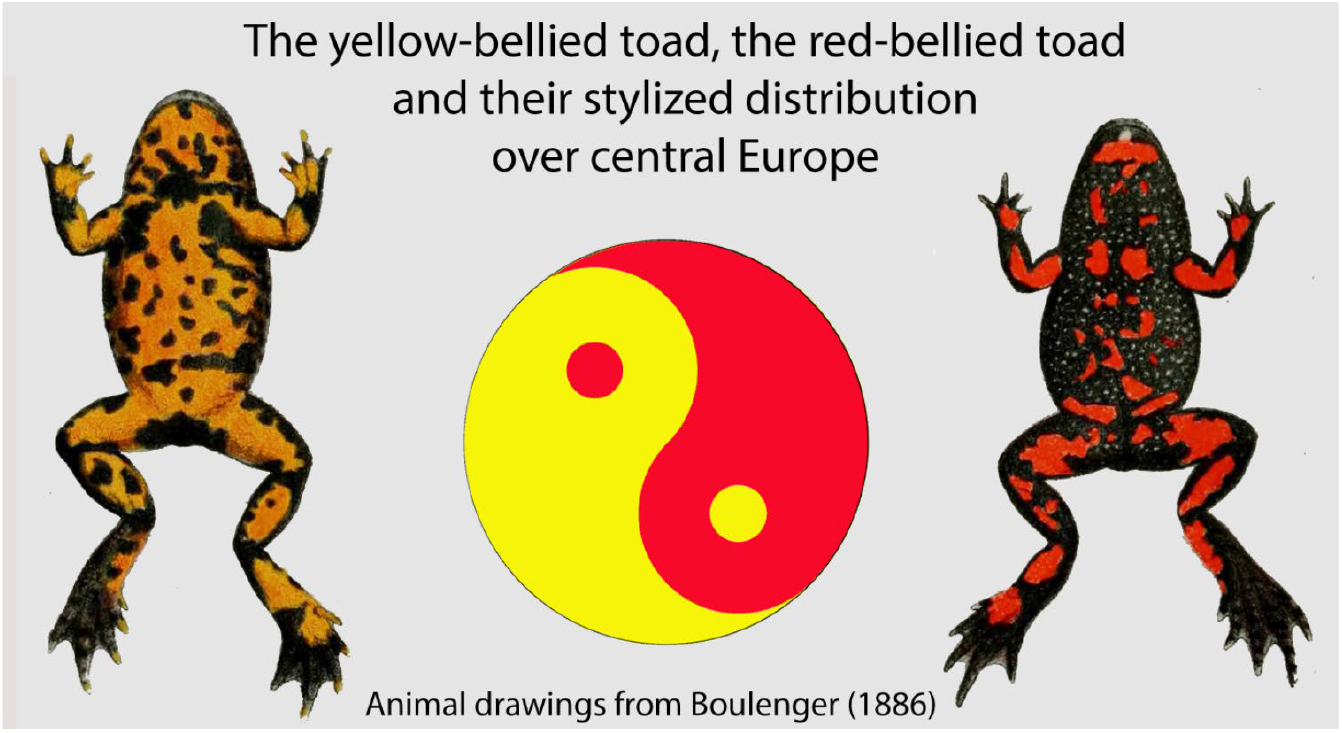

**Highlights:**

- Red- and yellow-bellied toads engage in a yin-yang-like mosaic distribution across Europe.
- Species’ differential habitat characteristics were studied with principal coordinates.
- Species distribution modelling was with climate data and with landscape variables.
- Models produced with atlas data for Hungary were assessed over neighbouring countries.
- Transferred models with elevation and forestation performed better than climate-based ones.

## 1 Introduction

Species distribution models (SDMs) are numerical tools that combine observations of species occurrence or abundance with environmental estimates with the goal to describe distributions across landscapes and to better understand the relationships of species and their environment (Thuiller, 2004; Wiens and Graham, 2005; Elith and Leathwick, 2009; Peterson et al., 2011). In the last decades, interest in SDM has greatly increased, with applications in fields wide apart as biogeography, systematics, ecology, evolution and conservation (Graham et al., 2004; Kozak and Wiens, 2006; Rodríguez et al., 2007, Pasanis et al., 2024). Whereas some SDM exercises include many candidate explanatory variables, motivated by their ready availability and a belief that the modelling procedure will identify those that are important, others have argued for the use only of those that are ecologically relevant to the target species (Ashcroft et al., 2011; Brodie et al., 2020; Sillero et al., 2021; Naas et al., 2024), so that the suitability estimated by the model reflects the species’ biology (Austin, 2002; Peterson and Nakazawa, 2008; Da Re et al., 2022). As was noted early on, the variable selection process and the inferences will improve if model building is based upon insight knowledge and insight (Mac Nally, 2002; Dormann, 2007), but unfortunately this premise is rarely assessed (Petitpierre et al., 2017; Regos et al., 2019; Lee-Yaw et al., 2022). While testing SDM forecasts is problematic because of the lack of reference data, ‘hindcasted’ models may be open for evaluation e.g., by paleopalynology for groups with extensive (sub)fossil data. A more promising avenue of research though is the spatial calibration of contemporary SDMs, with models built in one region and predictions evaluated in other regions (Blois et al., 2013).

We here present a case study on model transferability to which we use publicly available data on the distribution of two competing toad species (*Bombina bombina* and *B. variegata*) in central Europe. Aim of the study is to test if explanatory variables selected from insight and knowledge (Arntzen, 2025) yield better SDMs than parameters without this perceived quality. Models are derived with data for the centrally located country Hungary (Herpterkep, 2025) and applied to neighbouring countries. The unusual mosaic species distribution of European *Bombina* toads, reminiscent of the yin-yan symbol (Figure 1), provides ample opportunity for the critical evaluation of transferred models.

**Figure 1.**
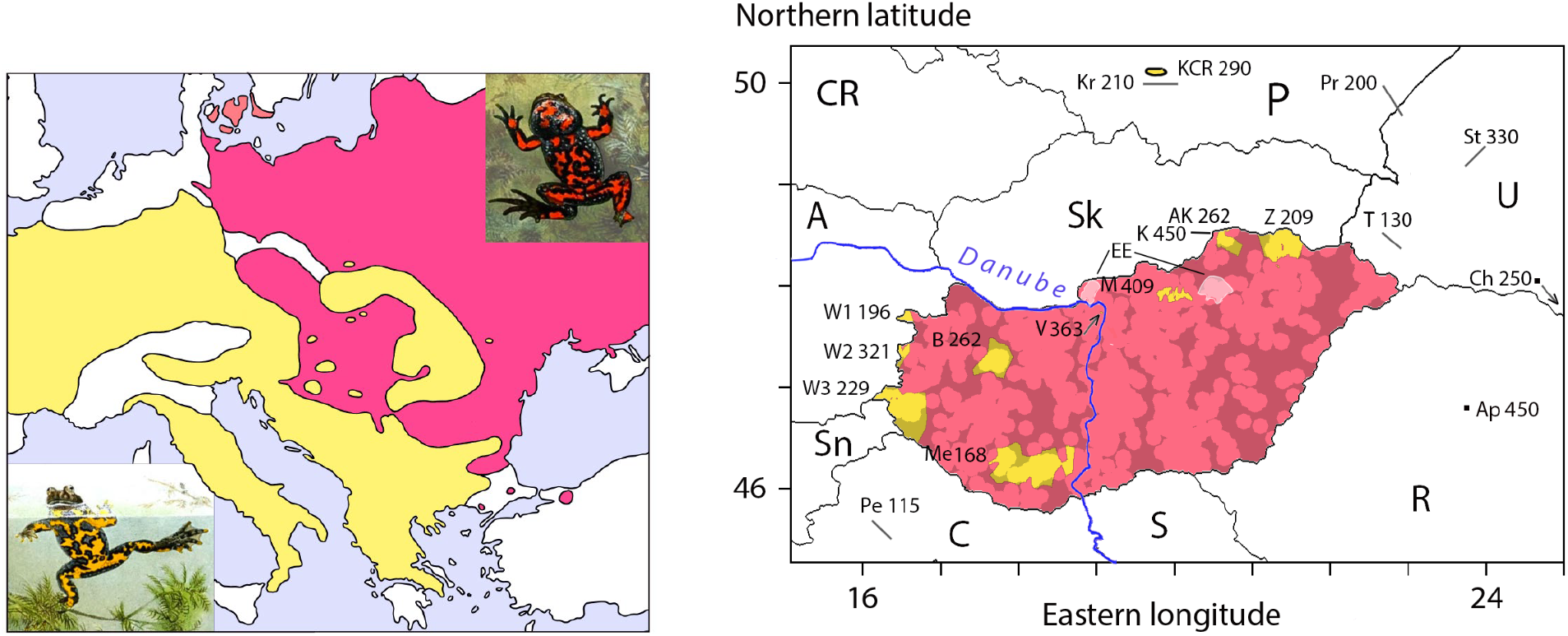
Distribution of *Bombina* species over central Europe. Panel left – range maps for the red-bellied toad *Bombina bombina* (in red) and the yellow-bellied toad *B. variegata* (in yellow) after Arntzen (1978) and Szymura (1993). Animal pictures are from Boulenger (1897) with *B. bombina* top right and *B. variegata* bottom left (see also Figure 5). Panel right – the distribution of *Bombina bombina* (in red) and *B. variegata* (in yellow) over Hungary from atlas data (Herpterkep, 2025) shown by a Dirichlet tessellation. Areas at distances of ca. 10 km or more from a recorded locality are grey shaded. Neighbouring countries are P – Poland, U – Ukraine, R – Romania, S – Serbia, C – Croatia, Sn – Slovenia, A – Austria, CR – Czech Republic and Sk – Slovakia. *Bombina variegata* areas in Hungary are, in clockwise order V – Visegrád mountains, M – Mátra mountains, AK – Aggtelek karst, Z – Zemplén mountains, Me – Mecsek mountains, B – Bakony mountains and W1-3 in the west of the country. KCR is a *B. variegata* enclave at the Kraków - Chrzanów ridge in southern Poland (Michałowski, 1961). Two white-shaded area in the north of the country – Börzsöny mountains to the west and Bükk mountains to the east – are ‘empty enclaves’ (EE), i.e., areas where experts would predict the occurrence of *B. variegata* but so far none were found (details see text). Lines indicate the centre positions of clinal *B. bombina – B. variegata* hybrid zones at K – karst area at the Slovakian – Hungarian border (Gollmann et al., 1988), Kr – Kraków (Szymura and Barton, 1986), Pr – Przemyśl (Pr, Szymura and Barton, 1991), T – Transcarpathian (Dufresnes et al., 2021), St – Stryi (Yanchukov et al., 2006) and Pe – PeŠćenica (MacCallum et al., 1998). Dots indicate hybrid belts in mosaic hybrid zones at Ch – Chernivtsi (Dufresnes et al., 2021) and A – Aphadia (Vines et al., 2003). Numbers indicate elevation at the species border, at the centre of the cline and at the hybrid belt, respectively. Other *Bombina*-hybrid zones known from the literature are not evaluated for sparse sampling or the absence of genetic data.

## 2 Material and methods

### 2.1 Research area and species distribution data

The area of research is central Europe with the focus on Hungary. Species distribution data from an atlas about to be published (Herpterkep, 2025) were used to *construct* a two-species distribution model (TSDM, see below). Independent data for *testing* TSDM performance were taken from the literature (Dufresnes et al., 2021) over the sector of *Bombina* species range overlap, bounded by the 14-28 eastern longitude and 43-51 northern latitude coordinates. Records not considered are duplications, interspecific hybrids and species syntopic occurrences at the pixel scale (see below). Excluded for model evaluation were records for Ukraine and Moldavia for which countries Corine land cover data are not available and Hungary for which the data may not be independent.

### 2.2 Environmental data

Environmental data considered as candidate explanatory variables to the reciprocal distribution of *B. bombina* and *B. variegata* were 19 climate variables (bio01–19) extracted from the WorldClim global climate database v.2 (Fick and Hijmans, 2017) (available at https://www.worldclim.org/data/index.html; for brief variable descriptions see Supplementary Information 1). For elevation we used the Copernicus digital elevation model (European Space Agency, 2024 available at https://doi.org/10.5069/G9028PQB). ‘Hilliness’ is the standard deviation of elevation derived with a 9*9-pixelwide (ca. 0.7 ha) filter. Vegetation data were from the Corine land cover database of the European Environment Agency (https://land.copernicus.eu/pan-european/corine-land-cover, in particular https://doi.org/10.2909/71fc9d1b-479f-4da1-aa66-662a2fff2cf7) (Büttner et al., 2021). Data were grouped in three layers as forestation (Corine classes 2, 3 and 4), shrub (class 5) and herbaceous (classes 6 and 7). The nominal resolution of the data is 30 arc-seconds for climate, 30 m for elevation and 10 m for land cover. An *a priori* distinction was made between variables that operate locally such as elevation, hilliness and land cover and those that take effect at larger spatial scale (i.e., the climate variables). To identify and subsequently reduce collinearity among the environmental variables, a half-matrix of the pairwise absolute Spearman correlation coefficients (r_S_) was subjected to clustering using the unweighted pair group with arithmetic mean method in Primer 7 (Dormann et al., 2013; Clarke and Gorley, 2015). Variables were retained using criteria of partial independence at |*r*_S_|<0.8 and were selected in alphanumerical order (Supplementary Information 1).

### 2.3 Canonical analysis of principal coordinates

The biological - environmental data set was analysed by Canonical Analysis of Principal coordinates (CAP) with Primer 7 and Permanova+ software, following the manuals (Anderson et al., 2008; Clarke and Gorley, 2015). Aim of the analysis is to find axes through the multivariate cloud of points that are best at discriminating the various groups of species occurrences. Groups considered were *B. bombina* (N=2310 randomly reduced to N=400 to limit the analytical workload), *B. variegata* (N=329) and four well-delimited *B. variegata* enclaves, namely the Bakony (N=21), Mátra (N=36), Mecsek (N=70) and Visegrád mountains (N=1) (see Figure 1A). Nineteen records that together describe a fifth well-resolved *B. variegata* enclave at the Kraków-Chrzanóv Ridge (KCR) in southern Poland are from Michałowski (1961). The Hungarian Börzsöny and Bükk mountains where *B. variegata* has not been observed but that in experts’ opinion (JV and KH) provide suitable habitat were included in the analysis by 50 randomly selected data points, for *a posteriori* consideration.

### 2.4 Two-species distribution modelling

Two-species distribution models in which the presence of one species is contrasted with the presence of the counterpart species (see Arntzen, 2023a) were derived with stepwise logistic regression analysis in SPSS v.30 (IBM SPSS, 2024). Parameter selection was in the forward stepwise mode under criteria of entry (P_in_=0.05) and removal of terms (P_out_=0.10) under the likelihood ratio criterion, while applying a weighing procedure that sets the number of records for species at par. Spatial models were analysed and visualized with ILWIS v.3.8.6 (ILWIS, 2019). The fit of the model to the underlying data was assessed by the area under the curve (AUC) statistic, as the other statistical evaluations done with SPSS 30 (IBM SPSS, 2024). Considering the possibility that hilliness better captures the overall habitat differentiation of *B. bombina* and *B. variegata* than elevation, models were rerun accordingly.

## 3 Results

The Hungarian atlas data show that the distribution of *B. bombina* and *B. variegata* is strongly parapatric, confirming earlier reconstructions (Figure 1). Four areas are resolved where *B. variegata* occurrences appear isolated from the continuous range by the counterpart species (i.e., enclaves), namely at the Bakony, Mátra, Mecsek and Visegrád mountains. Elevations coinciding with the range border vary from 168 to 409 m. At another five Hungarian areas that form part of the main *B. variegata* range, elevations at the range border vary from 196 to 431 m. Finally, data from the hybrid zone literature for neighbouring countries show species transitions at elevations varying from 115 to 450 m (Figure 1B). For the Kraków transect the elevation coinciding with the centre of the hybrid zone is 210 m, in line with an average elevation of 290 m at the range border at the adjacent KCR enclave. At the two ‘empty enclaves’ (i.e., without *B. variegata*) *B. bombina* records range up to 574 m in the Bükk mountains (Dely, 1966) and up to 747 m at Börzsöny (Herpterkep, 2025). High-elevation occurrences of *B. bombina* in the nearby presence of *B. variegata* are found in Mátra (localities at 510 and 650 m, Szabó, 1959) and the the Zemplén mountains (510-600 m).

Environmental parameters selected for analysis were six climatic variables and the locally operating variables elevation, forestation and herbaceous vegetation cover (Supplementary Information 1). Canonical analysis of principal coordinates separates the two species to which the contributing parameters are forestation, elevation and three precipitation parameters (bio12, bio14 and bio15) that are positively associated with *B. variegata* versus five temperature parameters and herbaceous vegetation cover that are positively associated with *B. bombina* (Figure 2). Records for the KCR-enclave fall in two groups inside the *B. variegata* and *B. bombina* clusters, respectively. The four Hungarian *B. variegata* enclaves group together along the forestation and elevation axes adjacent to continuous *B. variegata*. Finally, the two empty enclaves are in part projected along the real enclaves and for another part within the *B. bombina* cluster.

**Figure 2.**
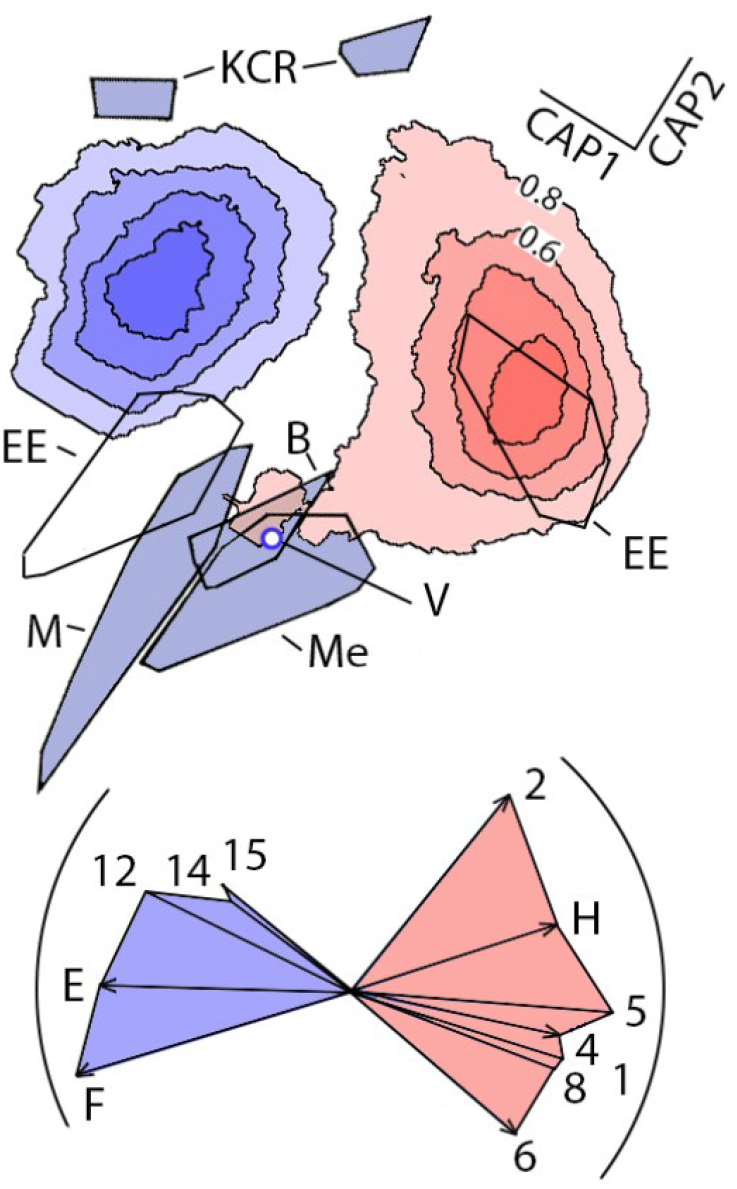
Top panel - bivariate plot of scores on the first and second axis analyses of a Canonical Analysis of Principal coordinates (CAP) with 12 selected environmental variables (see Supplementary Information 1) and various groups of *Bombina*-toads. Continuous range results are summarized by data density plots with *B. variegata* in blue and *B. bombina* in red, with increments of 0.2. Data for *B. variegata* enclaves are summarized by steel grey convex outline polygons and coded as: B – Bakony forest, KCR – the Kraków - Chrzanów ridge (in two sections), M – Mátra mountains, Me – Mecsek mountains and by a white dot for V – Visegrád mountains. Data for two ‘empty enclaves’ (EE, details see text) are pooled and yet consist of two, widely separated components associated with *B. variegata* and *B. bombina*, respectively. Bottom panel – arrows obtained by CAP indicate the impact (by length relative to the maximum shown by the partial circle) and the association with (by orientation) the listed parameters, as positively correlated with *B. variegata* records (blue shaded surface) and *B. bombina* (red shaded surface). Numbers refer to bioclimatic variables in which 1-8 are temperature related and 12-15 are precipitation related (for brief descriptions see Supplementary Information 1). E – elevation, H – herbaceous and F – forestation. Both panels are rotated by c. 30 degrees clockwise so that the horizontal axis is in direction of the parameter elevation that is traditionally used to capture the different ecologies of the species (Mertens, 1928; Arntzen, 1978; Dufresnes et al., 2021).

The TSDM for climatic variables is *P*=(1/(1+exp(4.194*bio01-4.471*bio05-1.008*bio06+0.432*bio08+0.0167*bio12+0.179*bio15+44.719))) and has a fit of AUC=0.983. The TSDM for local parameters is *P*=(1/(1+exp(0.00842*elevation+5.761*forestation-0.659*herbaceous-4.744))) with a model fit of AUC=0.954. For 95% confidence intervals of the estimates see Figure 3. Although the AUC curves are significantly different (ΔAUC=-0.029, Delong test z=7.405, P<0.0001), the projected species distributions are similar across Hungary, with as a notable exception the modelled occurrence of *B. variegata* at Lake Balaton (Figure 4). Evaluated against the reference data set of species presences outside Hungary (Supplementary Information 2), model fit is moderate for the climate parameters (AUC=0.792) and high for the local parameters (AUC=0.912), with model fit values that dropped by ΔAUC=0.191 (z=6.320, P<0.0001) and by ΔAUC=0.042 (z=2.107, P<0.05). The model with all 12 variables available for selection (representing a traditional research design) incorporates elevation, forestation and eight climate variables and performed well in Hungary (AUC=0.987) and less so abroad (AUC=0.801), also representing a significand drop (ΔAUC=0.186, z=6.061, P<0.0001). Finally, models were re-estimated with hilliness instead of elevation as a locally operating variable to which AUC model fit significantly dropped, in descriptive mode (from AUC=0.954 to 0.942, ΔAUC=0.012, z=7.151, P<0.0001) as well is in transference mode (from AUC=0.912 to 0.842, ΔAUC=0.070, P<0.0001) (Figure 3).

**Figure 3.**
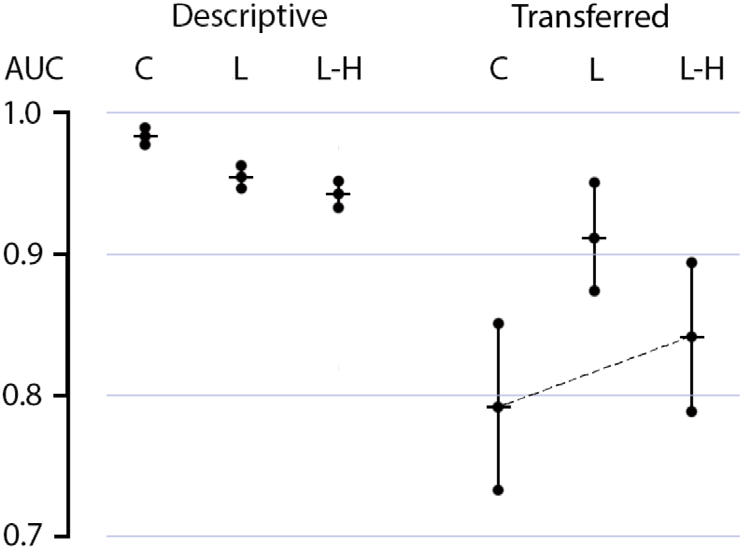
Fit of two-species distribution models for *Bombina bombina* and *B. variegata* from Hungarian atlas data (left – descriptive) and to reference data from abroad (right – upon spatial transference, see also Supplementary Information 2) described by the Area Under the Curve (AUC) statistic. Parameter sets from which the models are derived are climate (C), locally operating (L), or locally operating with hilliness taking the place of elevation (L-H). Vertical bars encompass the 95% confidence interval. Model fit values are significantly different among data sets (see main text), except those connected by an interrupted line (P<0.05).

**Figure 4.**
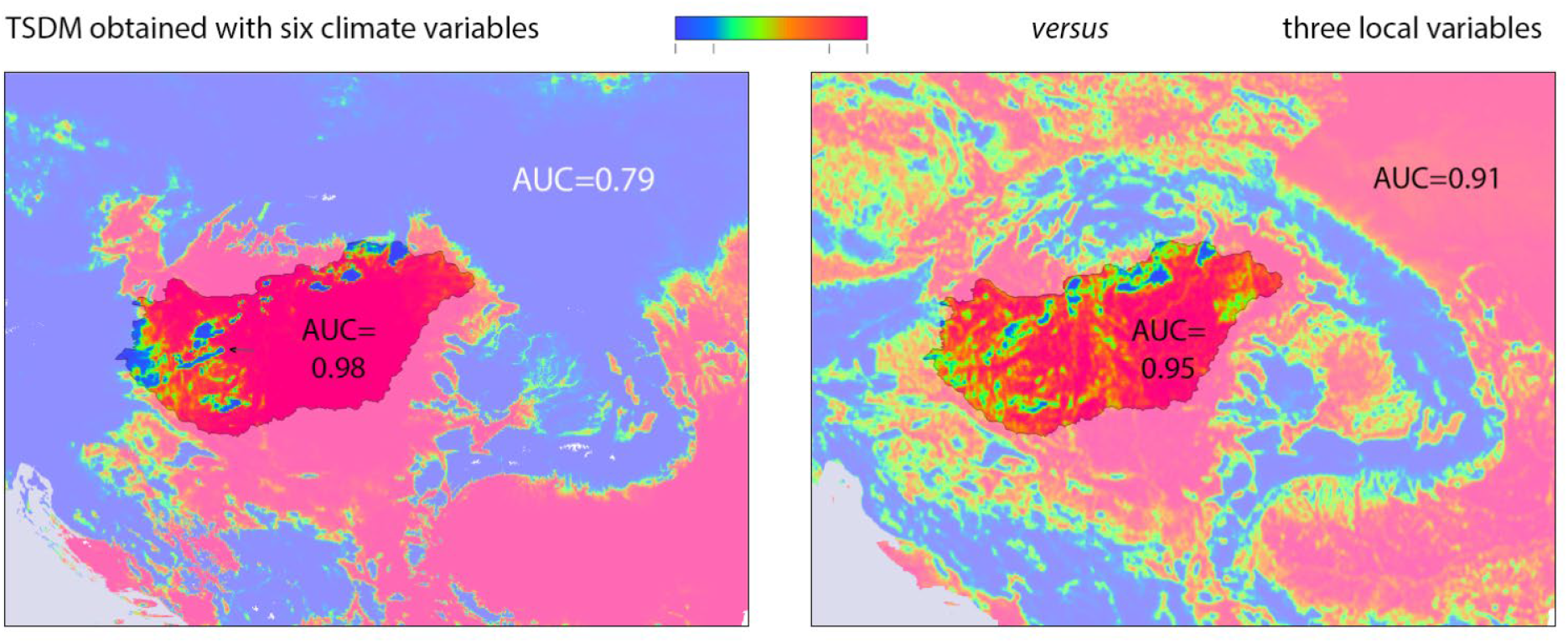
Two-species distribution models for *B. bombina* - *B. variegata* derived from Hungarian atlas data, with predicted areas of occurrence in red and blue. For intermediate probabilities of species occurrence in orange and green (0.2<*P*<0.8; see colour legend). An anomalous prediction of *B. variegata* at Lake Balaton is shown by an arrow. Note that, while model fits from six climate variables (left) versus three local parameters (right, smoothened for presentation purposes) are similar, the transferability of the ‘climate’ model is inferior to that of the ‘local’ model. The reference data with which the models are tested are shown in Supplementary Information 3.

## 4 Discussion

The fire-bellied toads *B. bombina* and *B. variegata* are distributed over several distinct and mutually exclusive areas across central Europe. At the interface of their essentially parapatric distributions the species engage in a long and winding contact zone where they also hybridize (Szymura, 1993) (Figure 1). Distributions cover mountainous and forested areas for *B. variegata* versus open lowlands for *B. bombina* (Arntzen, 2025). The principal difference between the species, however, resides not in landscape affiliation, but in breeding habitats that are typically small and ephemeral water bodies for *B. variegata* versus large and more permanent ponds for *B. bombina* (Figure 5). Accordingly, forestation and elevation (with hilliness as a possible substitute) are essential proxy parameters because obtaining blanket coverage of aquatic sites would involve an enormous effort. The critical issue here addressed is the relative performance in SDM of the locally operating landscape proxies versus the large-scale operating climate variables. The species’ intricate distribution mosaic is uniquely suited to test the usefulness of different types of data for SDM, because the species mosaic covers source and test areas alike. Using atlas data from the centrally located country Hungary we here extract the species’ ecological profiles with CAP, and we construct TSDMs that are subsequently tested for transferability with data from surrounding countries.

**Figure 5.**
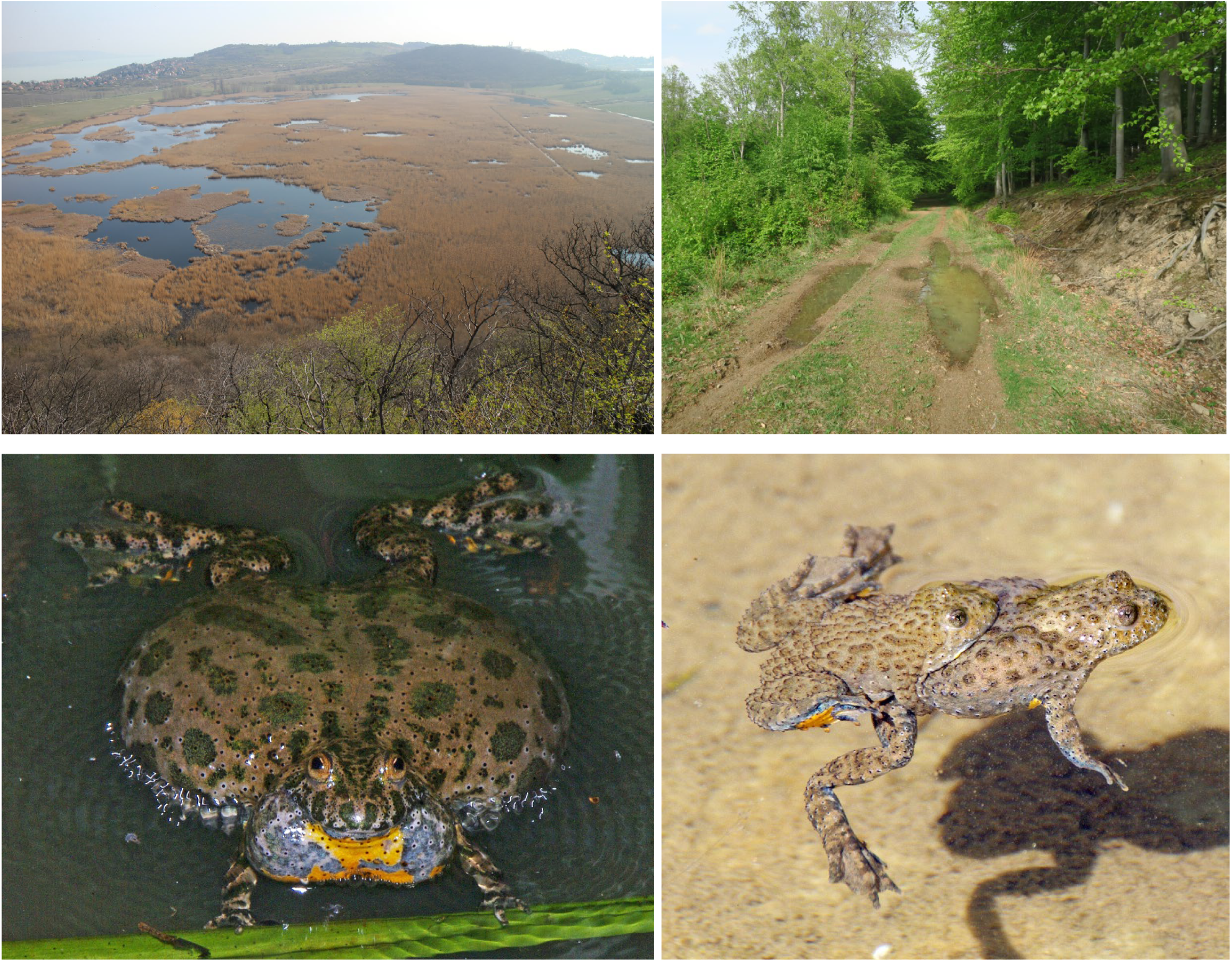
Top row - breeding localities of European *Bombina* toads with to the left the lake Külső-tó at Tihany peninsula, Hungary as typical for *B. bombina* and to the right temporary car track puddles in the Mátra mountains, Hungary as typical for *B. variegata*. Photos by JV. Bottom row – left, calling *B. bombina* male from Siemianówka, Poland and right an amplexed pair of *B. variegata* at the Saint Dionysios monastery, near Litochoro, Greece. Photo courtesy Serge Bogaerts.

The CAP-profiles are highly discriminatory with *B. bombina* associated to open lowland and higher temperatures and *B. variegata* associated with forested hills and mountains with higher precipitation (Figure 2). It is remarkable that the parameter with the largest impact on the separation of species with CAP is forestation and not, as traditionally recognized, elevation (Von Méhely, 1905; Mertens, 1928; Arntzen, 1978; Dufresnes et al., 2021). Four Hungarian *B. variegata* enclaves take similar positions in the bivariate plot and are more associated with elevation and forestation than with precipitation. The Polish KCR enclave contains more than the typical *B. variegata* habitat, possibly due to the inclusion of *B. bombina – B. variegata* hybrid populations along its fringes, or to a biased species identification from toad belly colouration characteristics where genetic data are yet unavailable (Michałowski, 1961). The position of the two ‘empty enclaves’ Börszöny and Bükk mountains in the CAP-plot suggests that prime *B. variegata* habitat is locally occupied by *B. bombina*, but also that the experts overestimated the ecological amplitude of *B. variegata*, as shown by the inclusion of *B. bombina* habitat, presumably representing the foothills rather than the mountain tops. Fossil information does not solve this issue. The Bükk mountains are karstic, with fossil-rich caves that support the presence of fire-bellied toads in the Pleistocene, but not what species (Jánossy, 1979) whereas the Börzsöny mountains have a volcanic origin and do not provide relevant information.

The TSDMs from climate data and from local variables both neatly describe the *Bombina* species mosaic in Hungary (Figures 3 and 4). However, the climate model is poorly transferable, whereas the model from the locally operating variables elevation and land cover performs well upon transference. Our results support the claim against the widespread practice of modelling with just climate data (Thuiller et al., 2004; Araújo and Rahbek, 2006; Dormann, 2007: Da Re et al., 2024, see e.g. Mi et al., 2024; Papežík et al., 2025). Whereas the inclusion of edaphic variables (e.g., ESDA European soil database v2.0, available at https://esdac.jrc.ec.europa.eu) may seem promising for terrestrial species that are in close connection to soil, such as many amphibians, this is demanding due to incomplete coverage, complicated data structures and lack of standardization (Arntzen et al., 2020). Conversely, land cover and elevation data are readily available, such as from the Corine and DEM databases here employed and otherwise from the ESA WorldCover project (https://esa-worldcover.org/en).

A weak tendency has been observed for a decline in SDM fit upon transference (Arntzen, 2006; Randin et al., 2006; Duncan et al., 2009; Rousseau and Betts, 2022; see Barbosa et al., 2009 for a counter example). However, the prime question is not how transferable the models are, but what explanatory parameters yield the best predictions as to improve our understanding of species-habitat associations. For European *Bombina* species these happen to be elevation and forestation, in line with knowledge and insights gathered earlier (Arntzen, 1978, 1996, 2025). Model fit is generally high which may be a consequence of the use of TSDM that is performed by *contrasting* species presences, so without inferred and difficult-to-justify absence data. The ‘biotic noise’ that is generally seen as a bias and a handicap in SDM (Elith and Leathwick, 2009; Anderson 2012; Sillero et al., 2021), is here turned to an advantage by focussing on the *differences* between parapatric species which is, however, at the expense of modelling results at the species’ distribution edges in allopatry (Arntzen, 2023a).

The European fire-bellied toads serve as an intriguing example but are not a one-off. Other pairs of European amphibians in which elevation and forestation are paramount in explaining a parapatric species distribution are *Lissotriton montandoni – L. vulgaris* (Babik et al., 2005; Antunes et al., 2023) and *Triturus cristatus – T. marmoratus* (Arntzen, 2023b). It is, however, the compounded, yin- and-yang symbol-like distribution of European *Bombina* species that allows a critical test of model transferability and evaluation of parameter performance. Although the TSDM of European fire-bellied toads from local variables performs well, it must be realized that that the variables are local indeed. The elevation at which *Bombina* species are separated in the TSDM is 220 m, but this is an omnibus value. Confined local models will identify other critical elevations, with elevation presumably fine-tuned somewhere within the 115-450 m range (Figure 1B). Local model adjustments from forestation and other land uses may similarly apply (Bugter et al., 1997; MacCallum et al., 1998; Arntzen, 2025 submitted). While there is just an indirect mechanistic relationship with the species’ differentiated niches, elevation and forestation are useful proxy variables that go a long way in capturing the more terrestrial mode of life of *B. variegata*, a species that preferentially breeds in small ephemeral ponds, as different from the more aquatic *B. bombina* that seeks to breed in larger and more permanent stagnant water bodies (Figure 5). These habitats are especially found in rolling, hilly and mountainous terrain for *B. variegata* versus flat areas for *B. bombina*. This basic habitat distinction was thought to perhaps be better captured by hilliness than by elevation (Arntzen, 1996), but this notion is not supported by the present study. The bottom-line messages are that even SDMs with near-perfect fit may perform poorly when applied outside areas for which they were designed and that an informed parameters selection enhanced model transferability, therewith improving our understanding of species-habitat associations.

## CRediT authorship contribution statement

Conceptualization – JWA; data curation – JWA, KH, JV; formal analysis – JWA; methodology – JWA; visualization – JWA; writing – original draft – JWA; writing – review and editing - JWA, JV.

## Data availability

This article does not contain new data. Occurrence data of *B. bombina* and *B. variegata* for Hungary are available at herpterkep.mme.hu. More precise coordinates than shown on the website may be provided upon request addressed to the Amphibian and Reptile Conservation Group of MME Birdlife Hungary at.

## References

Anderson, M. J., Gorley, R. N., Clarke, K. E. (2008) Permanova+ for Primer. Guide to Software and Statistical Methods. PRIMER-E, Plymouth, United Kingdom.

Anderson, R. P. (2012) Harnessing the world’s biodiversity data: promise and peril in ecological niche modeling of species distributions. Annals of the New York Academy of Sciences 1260: 66–80. 10.1111/j.1749-6632.2011.06440.x.

Antunes, B., Figueiredo-Vázquez, C., Dudek, K., Liana, M., Pabijan, M., Zieliński, P., Babik, W. (2023) Landscape genetics reveals contrasting patterns of connectivity in two newt species (Lissotriton montandoni and L. vulgaris). Molecular Ecology 32: 4515–4530. 10.1111/mec.16543

Araujo, M. B., Rahbek, C. (2006) How does climate change affect biodiversity? Science 313(5792): 1396–1397. 10.1126/science.1131758

Arntzen, J. W. (1978) Some hypotheses on postglacial migrations of the fire-bellied toad, Bombina bombina (Linnaeus) and the yellow-bellied toad, Bombina variegata (Linnaeus). Journal of Biogeography 5: 339–345. 10.2307/303802

Arntzen, J. W. (1996) Parameters of ecology and scale integrate the gradient and mosaic models of hybrid zone structure in Bombina toads and Triturus newts. Israel Journal of Ecology and Evolution 42: 111–119.

Arntzen, J. W. (2006) From descriptive to predictive distribution models: a working example with Iberian amphibians and reptiles. Frontiers in Zoology 3: 1–11. 10.1186/1742-9994-3-8

Arntzen, J. W. (2023a) A two-species distribution model for parapatric newts, with inferences on their history of spatial replacement. Biological Journal of the Linnean Society 138: 75–88. 10.1093/biolinnean/blac134

Arntzen, J. W. (2023b) Patch analysis of atlas data reveals pattern and process of species replacement. Frontiers of Biogeography 15: 3. 10.21425/F5FBG59627

Arntzen, J. W. (2025) Long term hybrid zone dynamics in red- and yellow-bellied toads estimated from environmental data at allopatric and parapatric scales. Manuscript submitted at preprint server

Arntzen, J. W., Canestrelli, D., Martínez-Solano, I. (2020) Environmental correlates of the European common toad hybrid zone. Contributions to Zoology 89: 270–281. 10.1163/18759866-bja10001

Ashcroft, M. B., French, K. O., Chisholm, L. A. (2011) An evaluation of environmental factors affecting species distributions. Ecological Modelling 222: 524–531. 10.1016/j.ecolmodel.2010.10.003

Austin, M. P. (2002) Spatial prediction of species distribution: an interface between ecological theory and statistical modelling. Ecological Modelling 157: 101–118. 10.1016/S0304-3800(02)00205-3

Babik, W., Branicki, W., Crnobrnja-Isailović, J., Cogălniceanu, D., Sas, I., Olgun, K., Poyarkov, N. A., Garcia-París, M., Arntzen, J. W. (2005) Phylogeography of two European newt species— discordance between mtDNA and morphology. Molecular Ecology 14: 2475–2491. 10.1111/j.1365-294X.2005.02605.x

Barbosa, A. M., Real, R., Vargas, M. J. (2009) Transferability of environmental favourability models in geographic space: The case of the Iberian desman (Galemys pyrenaicus) in Portugal and Spain. Ecological Modelling 220: 747–754. 10.1016/j.ecolmodel.2008.12.004

Blois, J. L., Williams, J. W., Fitzpatrick, M. C., Jackson, S. T., Ferrier, S. (2013) Space can substitute for time in predicting climate-change effects on biodiversity. Proceedings of the National Academy of Sciences 110: 9374–9379. 10.1073/pnas.1220228110

Boulenger GA (1886) On two European species of Bombinator. Proceedings of the Scientific Meetings of the Zoological Society of London 1886: 499–501. https://www.biodiversitylibrary.org/page/30829774#page/624/mode/1up

Boulenger, G. A. (1897) The Tailless Batrachians of Europe. Ray Society. London, United Kingdom. 10.5962/bhl.title.57744

Brodie, S. J., Thorson, J. T., Carroll, G., Hazen, E. L., Bograd, S., Haltuch, M. A., Holsman, K. K., Kotwicki, S., Samhouri, J. F., Willis-Norton, E., Selden, R. L. (2020) Trade-offs in covariate selection for species distribution models: a methodological comparison. Ecography 43: 11–24. 10.1111/ecog.04707

Bugter, R., MacCallum, C. J., Arntzen, J. W., Barton, N. H. (1997) An analysis of habitat components in a Bombina hybrids zone using remote sensing. Pp. 35-42 in Herpetologia Bonnensis (W. Böhme, W. Bischoff, T. Ziegler, eds). Societas Europaea Herpetologica, Bonn, Germany.

Büttner, G., Kosztra, B., Maucha, G., Pataki, R., Kleeschulte, S., Hazeu, G. W., Vittek, M., Schroder, C., Littkopf, A. (2021). Copernicus Land Monitoring Service – CORINE Land Cover. User Manual. Copernicus Publications. Göttingen, Germany.

Clarke, K. R., Gorley, R. N. (2015) PRIMER v7: User Manual/Tutorial. PRIMER-E, Plymouth, United Kingdom.

Da Re, D., Tordoni, E., Lenoir, J., Rubin, S., Vanwambeke, S. O. (2024) Towards causal relationships for modelling species distribution. Journal of Biogeography 51: 840–852. 10.1111/jbi.14775

Dely, G. O. (1966) Amphibians and reptiles of the Bükk Mountains. Pp. 535–570 in: Mahunka, S. (ed.): The fauna of the Bükk National Park. Hungarian Natural History Museum, Budapest, Hungary.

Dormann, C. F. (2007) Promising the future? Global change projections of species distributions. Basic and Applied Ecology 8: 387–397. 10.1016/j.baae.2006.11.001

Dormann, C. F., Elith, J., Bacher, S., Buchmann, C., Carl, G., Carré, G., García Marquéz, J. R., Gruber, B., Lafourcade, B., Leitão, P., Münkemüller, T., McClean, C., Osborne, P. E., Reineking, B., Schröder, B., Skidmore, A. K., Zurell, D.,Lautenbach, S. (2013) Collinearity: a review of methods to deal with it and a simulation study evaluating their performance. Ecography 36: 27–46. 10.1111/j.1600-0587.2012.07348.x

Dufresnes, C., Suchan, T., Smirnov, N. A., Denoël, M., Rosanov, J. M., Litvinchuk, S. N. (2021) Revisiting a speciation classic: Comparative analyses support sharp but leaky transitions between Bombina toads. Journal of Biogeography 48: 548–560. 10.1111/jbi.14018

Duncan, R. P., Cassey, P., and Blackburn, T. M. (2009) Do climate envelope models transfer? A manipulative test using dung beetle introductions. Proceedings of the Royal Society B: Biological Sciences 276: 1449–1457. 10.1098/rspb.2008.1801

Elith, J., Leathwick, J. R. (2009) Species distribution models: ecological explanation and prediction across space and time. Annual Review of Ecology, Evolution, and Systematics 40: 677–697. 10.1146/annurev.ecolsys.110308.120159

European Space Agency (2024) Copernicus Global Digital Elevation Model. Distributed by OpenTopography. Accessible at 10.5069/G9028PQB

Fick, S. E., Hijmans, R. J. (2017) WorldClim 2: new 1-km spatial resolution climate surfaces for global land areas. International Journal of Climatology 37: 4302–4315. 10.1002/joc.5086

Gollmann, G., Roth, P., Hödl, W. (1988) Hybridization between the fire-bellied toads Bombina bombina and Bombina variegata in the karst regions of Slovakia and Hungary: morphological and allozyme evidence. Journal of Evolutionary Biology 1: 3–14. 10.1046/j.1420-9101.1988.1010003.x

Graham, C. H., Ron, S. R., Santos, J. C., Schneider, C. J., Moritz, C. (2004) Integrating phylogenetics and environmental niche models to explore speciation mechanisms in dendrobatid frogs. Evolution 58: 1781–1793. 10.1111/j.0014-3820.2004.tb00461.x

Herpterkep (2025) Amphibian and Reptile Mapping of the Amphibian and Reptile Conservation group. MME Birdlife Hungary (2025) Accessible at herpterkep.mme.hu

IBM Corp. (2024) IBM SPSS Statistics for Windows, Version 30.0. IBM Corp, Armonk, New York, USA.

ILWIS (2019) Integrated Land and Watershed Management Information System. International Institute for Aerospace Survey and Earth Sciences, Enschede, The Netherlands.

Jánossy D. (1979) Classification of the Pleistocene in Hungary based on the vertebrate fauna [In Hungarian]. Akadémiai Kiadó, Budapest, Hungary.

Kozak, K. H., Wiens, J. (2006) Does niche conservatism promote speciation? A case study in North American salamanders. Evolution 60: 2604–2621. 10.1111/j.0014-3820.2006.tb01893.x

Lee-Yaw, J. A., McCune, J. L., Pironon, S., Sheth, S. N. (2022) Species distribution models rarely predict the biology of real populations. Ecography 2022: e05877. 10.1111/ecog.05877

Mac Nally, R. (2002) Multiple regression and inference in ecology and conservation biology: further comments on identifying important predictor variables. Biodiversity and Conservation 11: 1397–1401. 10.1023/A:1016250716679

MacCallum, C. J., Nürnberger, B., Barton, N. H., Szymura, J. M. (1998) Habitat preference in the Bombina hybrid zone in Croatia. Evolution 52: 227–239. 10.2307/2410938

Mertens, R. (1928) Zur Naturgeschichte der europäishen Unken (Bombina). Zeitschrift für Morphologie und Ökologie der Tiere 11: 613–633. 10.1007/BF02424588

Mi, C., Han, X., Jiang, Z., Zeng, Z., Du, W., Sun, B. (2024) Precipitation and temperature primarily determine the reptile distributions in China. Ecography 2024: e07005. 10.1111/ecog.07005

Michałowski, J. (1961) The Bombinators of the Kraków-Chrzanów Ridge and the adjacent part of the river Wisła valley. Acta Zoologica Cracoviensia 5: 699–714.

Naas, A. E., Keetz, L. T., Halvorsen, R., Horvath, P., Mienna, I. M., Simensen, T., Bryn, A. (2024) Choice of predictors and complexity for ecosystem distribution models: effects on performance and transferability. Ecography 2024: e07269. 10.1111/ecog.07269

Papežík, P., S Aschengeschwandtnerová, S., Benovics, M., Dedukh, D., Doležálková-Kaštánková, M., Choleva, L., Javorčík, A., Lymberakis, P., Papežíková, S., Poulakakis, N., Sarı, İ., Šanda, R., Vukić, J., Mikulíček, P. (2025) A pattern of hybridisation and population genetic structure of two water frog species (Ranidae, Amphibia) in the southwestern Balkans. Zoologica Scripta xx: 1–22. 10.1111/zsc.12723

Pasanisi, E., Pace, D. S., Orasi, A., Vitale, M., & Arcangeli, A. (2024) A global systematic review of species distribution modelling approaches for cetaceans and sea turtles. Ecological Informatics 2024: 102700. 10.1016/j.ecoinf.2024.102700

Peterson, A. T., Nakazawa, Y. (2008) Environmental data sets matter in ecological niche modelling: an example with Solenopsis invicta and Solenopsis richteri. Global Ecology and Biogeography 17: 135–144. 10.1111/j.1466-8238.2007.00347.x

Peterson, A. T., Soberón, J., Pearson, R. G., Anderson, R. P., Martínez-Meyer, E., Nakamura, M., Araújo, M. B. (2011) Ecological Niches and Geographic Distributions. Princeton University Press, Princeton, USA. 10.23943/princeton/9780691136868.003.0003

Petitpierre, B., Broennimann, O., Kueffer, C., Daehler, C., Guisan, A. (2017) Selecting predictors to maximize the transferability of species distribution models: Lessons from cross-continental plant invasions. Global Ecology and Biogeography 26: 275–287. 10.1111/geb.12530

Randin, C. F., Dirnböck, T., Dullinger, S., Zimmermann, N. E., Zappa, M., Guisan, A. (2006) Are niche-based species distribution models transferable in space? Journal of Biogeography 33: 1689–1703. 10.1111/j.1365-2699.2006.01466.x

Regos, A., Gagne, L., Alcaraz-Segura, D., Honrado, J. P., Domínguez, J. (2019) Effects of species traits and environmental predictors on performance and transferability of ecological niche models. Scientific Reports 9: 4221. 10.1038/s41598-019-40766-5

Rodríguez, J. P., Brotons, L., Bustamante, J., Seoane, J. (2007) The application of predictive modelling of species distribution to biodiversity conservation. Diversity and Distributions 13: 243–251. 10.1111/j.1472-4642.2007.00356.x

Rousseau, J. S., Betts, M. G. (2022) Factors influencing transferability in species distribution models. Ecography 2022: e06060. 10.1111/ecog.06060

Sillero, N., Arenas-Castro, S., Enriquez-Urzelai, U., Vale, C. G., Sousa-Guedes, D., Martínez-Freiría, F., Real, R., Barbosa, A. M. (2021) Want to model a species niche? A step-by-step guideline on correlative ecological niche modelling. Ecological Modelling 456: 109671. 10.1016/j.ecolmodel.2021.109671

Szabó, I. (1959) Contributions à la répartition du Sonneur aux pieds épais (Bombina variegata Linné) en Hongrie. Vertebrata Hungarica 1: 161–169.

Szymura, J. M. (1993) Analysis of hybrid zones with Bombina. Pp. 261-289 in: Hybrid Zones and the Evolutionary Process (R. G. Harrison, editor). Oxford University Press, United Kingdom. 10.1093/oso/9780195069174.003.0010

Szymura, J. M., Barton, N. H. (1986) Genetic analysis of a hybrid zone between the fire-bellied toads, Bombina bombina and B. variegata, near Cracow in southern Poland. Evolution 40: 1141–1159. 10.2307/2408943

Szymura, J. M., Barton, N. H. (1991) The genetic structure of the hybrid zone between the fire-bellied toads Bombina bombina and B. variegata: comparisons between transects and between loci. Evolution 45: 237–261. 10.2307/2409660

Thuiller, W., Lavorel, S., Midgley, G. U. Y., Lavergne, S., Rebelo, T. (2004) Relating plant traits and species distributions along bioclimatic gradients for 88 Leucadendron taxa. Ecology 85: 1688–1699. 10.1890/03-0148

Vines, T. H., Köhler, S. C., Thiel, M., Ghira, I., Sands, T. R., MacCallum, C. J., Barton, N. H., Nürnberger, B. (2003) The maintenance of reproductive isolation in a mosaic hybrid zone between the fire-bellied toads Bombina bombina and B. variegata. Evolution 57: 1876–1888. 10.1111/j.0014-3820.2003.tb00595.x

Von Méhély, L. (1905) Die herpetologischen Verhaltnisse des Mecsek-Gebirges und der Kapela. Annales historico-naturales Musei Nationalis Hungarici Budapest 3: 256–316.

Wiens, J. J., Graham, C. H. (2005) Niche conservatism: integrating evolution, ecology, and conservation biology. Annual Review of Ecology, Evolution and Systematics 36: 519–539. 10.1146/annurev.ecolsys.36.102803.095431

Yanchukov, A., Hofman, S., Szymura, J. M., Mezhzherin, S. V., Morozov-Leonov, S. Y., Barton, N. H., Nürnberger, B. (2006) Hybridization of Bombina bombina and B. variegata (Anura, Discoglossidae) at a sharp ecotone in western Ukraine: comparisons across transects and over time. Evolution 60: 583–600. 10.1554/04-739.1

