## Supplementary Information for "Assessment of environmental variables for species distribution modelling: insights from the mosaic distribution of red- and yellow-bellied toads"

**Supplementary Information 1**

Clustering of environmental data with the unweighted pair group method with arithmetic mean on basis of the absolute value of Spearman’s correlation coefficient at |r_S_|= 0.8. Variable selection was in alphanumerical order and land cover data for ‘shrub’ were not used for reason of low information content. Selected variables are indicated by a black dot. The climatic variables area: bio01 – annual mean temperature, bio02 – mean diurnal range (mean of monthly (max temp - min temp)), bio03 – isothermality (bio02/bio07) (×100), bio04 – temperature seasonality (standard deviation ×100), bio05 – maximum temperature of warmest month, bio06 – minimum temperature of coldest month, bio007 – temperature annual range (bio05-bio06), bio08 – mean temperature of wettest quarter, bio9 – mean temperature of driest quarter, bio10 – mean temperature of warmest quarter, bio11 – mean temperature of coldest quarter, bio12 – annual precipitation, bio13 – precipitation of wettest month, bio14 – precipitation of driest month, bio15 – precipitation seasonality (coefficient of variation), bio16 – precipitation of wettest quarter, bio17 – precipitation of driest quarter, bio18 – precipitation of warmest quarter and bio19 – precipitation of coldest quarter.

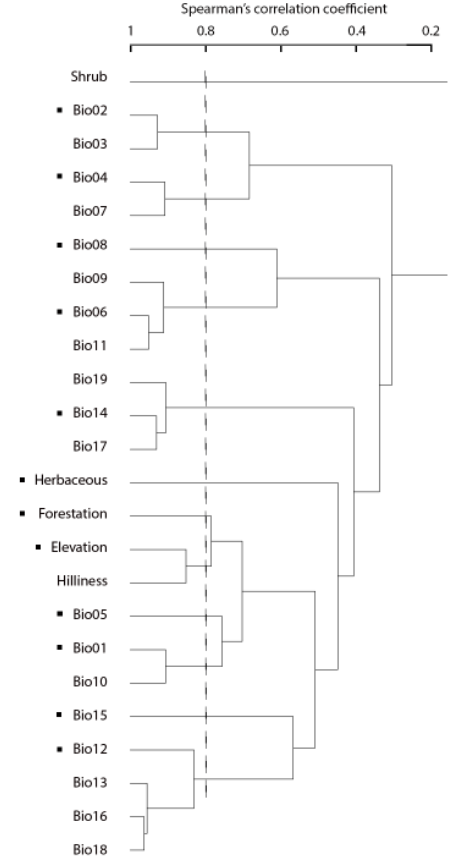

**Supplementary Information 2**

Records for the red-bellied toad *Bombina bombina* (in red) and the yellow-bellied toad *B. variegata* (in blue) from central Europe used to test two-species distribution models derived from Hungarian atlas data. The data are from Dufresnes et al. (2021). Records with a zero decimal coordinate precision are not shown. Data for Ukraine and Moldavia (shaded) were not considered for model testing due to the absence of land cover data for those countries. Data for Hungary (shaded) may not be independent and were also excluded. Citation: Dufresnes, C., Suchan, T., Smirnov, N. A., Denoël, M., Rosanov, J. M., Litvinchuk, S. N. (2021) Revisiting a speciation classic: Comparative analyses support sharp but leaky transitions between *Bombina* toads. *Journal of Biogeography 48*: 548-560.

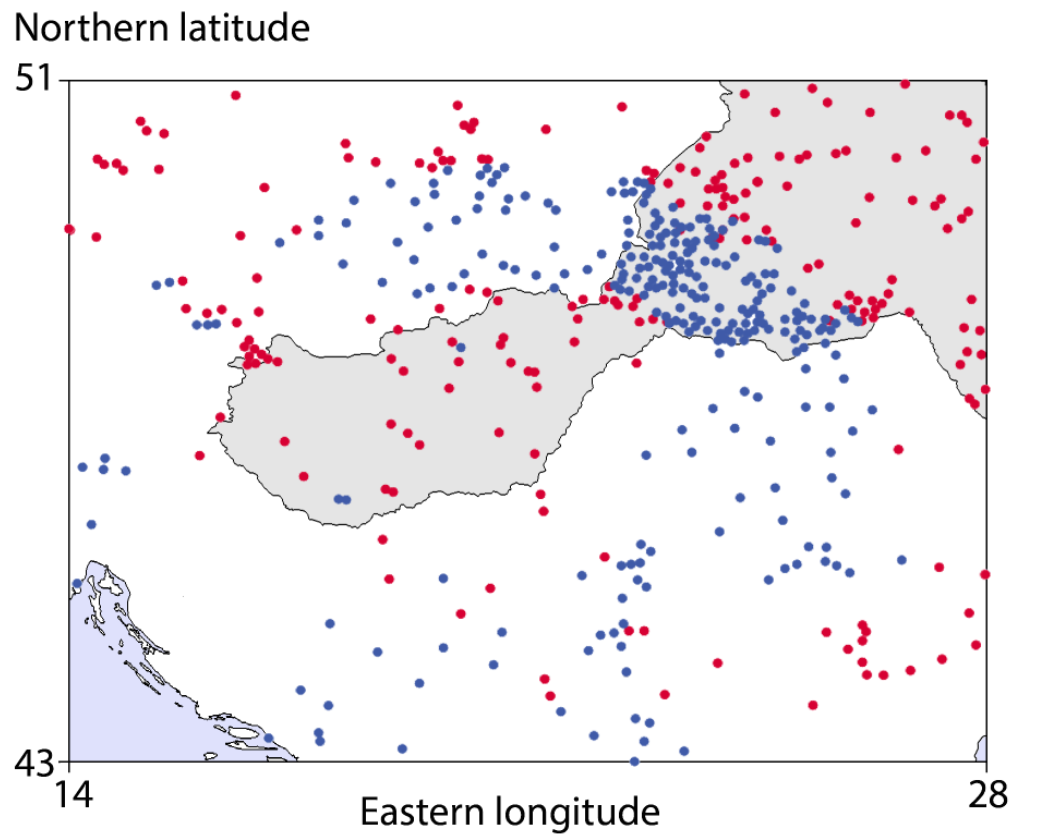
